# Intra-Specific variability in fluctuating environments - mechanisms of impact on species diversity

**DOI:** 10.1101/767491

**Authors:** Bnaya Steinmetz, Michael Kalyuzhny, Nadav M. Shnerb

**Affiliations:** Department of Physics, Bar-Ilan University, Ramat-Gan IL52900, Israel; Department of Ecology, Evolution, and Behavior, Institute of Life Sciences, Hebrew University of Jerusalem, Givat-Ram, Jerusalem, 91904, Israel

## Abstract

Recent studies have found considerable trait variations within species. The effect of such intra-specific trait variability (ITV) on the stability, coexistence and diversity of ecological communities received considerable attention and in many models it was shown to impede coexistence and decrease species diversity. Here we present a numerical study of the effect of genetically inherited ITV on species persistence and diversity in a temporally fluctuating environment. Two mechanisms are identified. First, ITV buffers populations against varying environmental conditions (portfolio effect) and reduces abundance variations. Second, the interplay between ITV and environmental variations tends to increase the mean fitness of diverse populations. The first mechanism promotes persistence and tends to increase species richness, while the second reduces the chance of a rare species population (which is usually homogenous) to invade and decreases species richness. We show that for large communities the portfolio effect is dominant, leading to ITV promoting species persistence and richness.

## I. INTRODUCTION

Understanding the factors and processes that shape the diversity of species in ecological communities is among the most important questions in ecology. Most current explanations of species diversity focus on the effects of niche differences and fitness-equalizing trade-offs [1–4], as well as spatial [5–7] and temporal [8–10] variability in environmental conditions. These factors affect the manner by which populations of different species interact and respond to the environment, and, consequently, determine their coexistence [2] and persistence [11]. However, until recently far less attention has been drawn to the possibility that the dynamics of a population may also be strongly affected by the variability of phenotypes exhibited by individuals within a population, or Intra-specific Trait Variability (ITV).

In recent years, ITV is drawing increasing attention [12–18]. This interest was triggered by the observation of considerable ITV in multiple ecological communities [14, 17], and motivated many studies of the mechanisms by which ITV influences ecological dynamics and biodiversity.

While considerable work has been done on the effect of ITV in deterministic settings, ecological dynamics also depends on stochastic events leading to temporal variations in abundance and possibly to the extinction of species that deterministically could exist [19]. In particular, demographic stochasticity refers to stochastic events acting on individuals independently of each other while environmental stochasticity, or environmental fluctuations, reflect erratic changes affecting entire populations synchronously. Environmental fluctuations have been shown to play a central role in shaping species abundances in multiple communities [20–23], and their effect on species diversity [24] and persistence [25–28] have been the focus of much contemporary work.

Here we would like to study the interplay between ITV and temporal environmental fluctuations, and to explore its effect on community dynamics and species richness. We will focus on the simplest (linear) form of environmental stochasticity, which affects reproduction and mortality simultaneously and does not generate any rare species advantage or disadvantage [9]. We consider heritable traits with different fitnesses, where a population supports a certain level of phenotypic diversity due to mutations. In such an eco-evolutionary setting the level of ITV is by itself a dynamical property that depends on the abundance of a population and responds to environmental variations.

Through this paper we identify and analyze two mechanisms by which the interaction between ITV and environmental stochasticity affects the dynamics of species competition. The first mechanism is the **portfolio effect**. In the presence of environmental fluctuations, the reproductive success of groups of individuals that share the same trait varies in synchrony, and the population abundance variance grows like the square of the group size. ITV breaks the synchrony between conspecific individuals, hence it reduces the strength of abundance variations for entire populations.

The portfolio effect was discussed a while ago in the context of fluctuations in the overall biomass of a community made of many species [29]. Its relevance to a single species with ITV (i.e., with many subspecies) was demonstrated in [30] and discussed in [12]. These authors focused their discussion on the effect of negative covariance between subspecies. Here we show that even when the covariance in the response of subspecies to environmental variations is zero, ITV yields a portfolio effect that modifies substantially the dynamic.

A more subtle mechanism has to do with the **adaptability advantage** of a more diverse population. In a fluctuating environment, at every instance of time intraspecific competition allows favoured sub-populations to increase in frequency, so the mean fitness of the entire population increases. Since ITV is correlated with population size (small populations tend to be homogenous), the adaptability advantage decreases the chance of rare populations to invade.

This mechanism may be considered as a specific example of the adaptive eco-evolutionary dynamic, i.e., the enhanced rate of adaptation caused by heritable trait variance [12, 31]. We will show that this effect exists even in a fluctuating environment, and even when the mean (over time) of the relative fitness of each trait is zero, provided that the correlation time of the environmental conditions, *τ*, allows for intraspecific competition to increase the frequency of the temporarily fitter subsepcies.

Interestingly, while both mechanisms tends to decrease the turnover rates, they have opposite effects on biodiversity. ITV diminishes abundance variations due to the portfolio effect, hence it increases coexistence time and allow a given system to support more diverse communities. On the other hand, low abundance species have lower chance to invade since they lack the adaptability advantage of the diverse. When rare populations cannot invade, extinction events are not compensated by establishment of new species and the overall species richness at equilibrium is reduced.

Through this paper we present the results of extensive numerical simulations and analyze them in light of previous theoretical studies. We consider the effect of ITV using three models with increasing levels of complexity:

- A two species competition model with complete niche overlap, each species is composed of many subspecies where the fitness of each subspecies is independent and its mean (over time) is zero (a time averaged neutral model as defined in [32]).
- A similar model that allows for speciation events, with a diverse steady state that reflects speciation-extinction equilibrium.
- A high-diversity, asymmetric, competitive Leslie-Gower model [33, 34] for annual plant community.

The first model is very simple and admits exact solutions without ITV, so it allows us to explore in detail the way in which the ITV portfolio effect reduces the impact of environmental stochasticity on abundance variations and makes them more and more demographic-like. In the second model the system supports many species and the interplay between the portfolio and the adaptability advantage effects is manifested explicitly in the species richness. The Leslie-Gower model allows us to examine our insights using a more realistic model that is widely used in the analysis of empirical data.

Our analysis suggests that in large communities the portfolio effect dominates over the adaptability advantage. Accordingly, in the models considered here the species richness of a community made of populations that support ITV is in general larger than the species richness of a similar community without ITV.

A few recent studies, in which environmental stochasticity was not taken into account, suggest that ITV would generally impede species coexistence and decrease richness [15, 16], mainly due to the increased degree of species interaction that leads to stronger effective competition. In the models we consider below ITV does not affect niche breadth, so this effect is absent. In empirical system ITV affects both niche breadth and the response to environmental fluctuations, so ITV may have stabilizing or destabilizing effects, depending on the relative strength of these mechanisms.

## II. THEORETICAL BACKGROUND

As explained above, when the environmental variations affects individuals in an uncorrelated manner the stochasticity of abundance fluctuations is demographic, and when the reproductive success of entire populations fluctuates coherently in time one speaks about environmental stochasticity. In a population that admits ITV subpopulations respond differently to the fluctuating environment, so this is an intermediate case between demographic and environmental stochasticity. In this section we provide a short survey of already known results about the persistence properties of communities under purely demographic stochasticity and under the combined effect of demographic and environmental stochasticity. We hypothesize that ITV will interpolate between these two extremes and use these known results as reference points. In stochastic models of species competition, many works examined how long it takes for diversity to be lost (in the absence of processes that reintroduce diversity, such as speciation). This can be quantified using the mean time (number of generations) to absorption *T*_*A*_(*x*), which is defined as the time until one of the species goes extinct, and essentially depends on the initial frequency *x*. In particular, if the dynamics of a two-species community, with community size of *J* individuals, is driven only by demographic stochasticity (a neutral model), it has been shown [35] that,

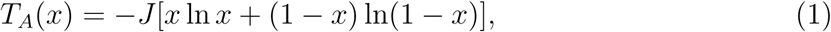

which implies that *T*_*A*_ scales linearly with community size *J*.

On the other hand, when the dynamic is still purely stochastic but the system is affected by both demographic and environmental stochasticity (a time-averaged neutral model), the mean time to absorption was found to satisfy [28],

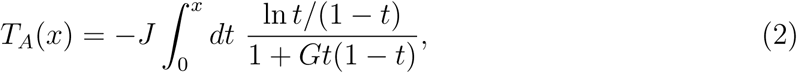

where *G* = *Jg* and *g* is the strength of the environmental stochasticity, as we shall explain below. When *J* is large the *J* dependence of this integral is given by,

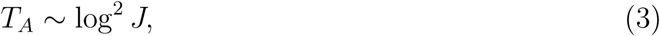

representing a dramatic decrease of persistence time due to the destabilizing effect of environmental stochasticity.

When *T*_*A*_ is low, extinction rates are high and, subsequently, species diversity at equilibrium is low. Hence, in competition models with environmental stochasticity which affects both mortality and reproduction, stochasticity tends to decrease species diversity by enhancing stochastic extinctions [24, 36].

## III. METHODS

We would like to investigate the consequences of ITV for population and community dynamics, species persistence, and eventually for species diversity, in stochastic, fluctuating environments. In particular, we would like to consider a case of competition between populations of different species, each exhibiting genetic variability, leading to variability in traits. We assume that the traits govern the response of individuals to environmental variations. As a result, a set of individuals within the species population having the same trait (referred to as sub-population) responds coherently to environmental fluctuations, while other sub-populations, having different traits, respond differently. We consider here only the effect of heritable trait variability on differential response to environmental variations.

We begin by presenting a simple, symmetric model of competition between species where mutations generate new traits (and therefore sub-populations), and new species may be formed by speciation. At any given moment, some traits (sub-populations) have higher fitness than the others. Accordingly, selection acts on the system in two levels: first, intraspecific selection tends to increase the relative fraction of high-fitness sub-populations within a given species population, then interspecific competition favors species whose mean fitness is high, i.e. populations dominated by individuals with favored traits. After some time, the environment changes and consequently the competitive abilities of all traits change. We analyze this model numerically and identify the two mechanisms mentioned above, namely the portfolio effect and the adaptability advantage of abundant populations.

We then study the combined effect of these mechanisms on the time to absorption *T*_*A*_(*x*) for a two species system, generally finding that the portfolio effect is dominant, making extinctions less frequent. We generalize these results to many species, examining neutral and non-neural multispecies models, again, finding that ITV promotes diversity by buffering population fluctuations.

### A. Models

#### 1. Symmetric models

To study the effect of ITV on species persistence and diversity, we will first analyze a symmetric, or Time-Averaged Neutral model of species competition. In such models, individuals compete according to their fitness, which depends on the environmental conditions. Environmental changes are modeled as temporal changes in fitness, whereas the fitnesses of all individuals are redrawn from the same distribution. Hence, while environmental fluctuations could change competitive hierarchy on the short run, over time these fluctuations average out and all individuals are fully symmetric. In previous symmetric models with environmental fluctuations [24, 32, 37], species respond coherently to the environment. Here, we allow some degree of genetic variability within each species, so two individuals of the same species may still have different *traits*. For simplicity, we will assume every individual has a single heritable trait (in addition to its species identity), and this trait governs its response to environmental fluctuations.

We study the dynamics of a continuous time process, having in mind a community of *J* competing individuals. To avoid nonlinear effects that may stabilize coexistence states via the storage mechanism [8], we use (as in [27, 38]) a simple model in which the relationships between fitness and growth rate are linear. We consider competition between two or more populations, where a chance encounter between two individuals may end up in a struggle over, say, a piece of food, a mate or a territory. To model that we pick, in each elementary birth-death event, two random individuals for a “duel”, the loser dies and the winner produces a single offspring. The chance to win is determined by the relative fitness: an individual with fitness *s*_1_ wins against an individual with *s*_2_ with probability,

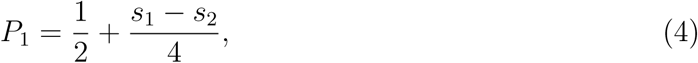

and the chance of the second to win is *P*_2_ = 1 − *P*_1_. With this “duel” competition, both mortality and reproduction are identically affected by fitness. This keeps the model linear [39], such that for two homogeneous species competition, where all individuals in species *i* have fitness *s*_*i*_, the mean frequency of species 1, *x*, satisfies the classical logistic equation

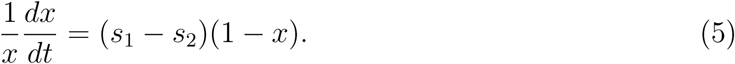

To introduce environmental fluctuations, we assume that fitness is fluctuating in time. Every individual’s fitness is determined by its trait *i*, resulting in the fitness of individuals with trait *i* at time *t* being *s*_*i*_(*t*). We have studied the simplest scenario of fitness fluctuations, in which *s*_*i*_(*t*) takes only two values (dichotomous stochasticity), either +*σ*_0_ or −*σ*_0_ (to keep 0 ≥ *P*_1_ ≤ 1, |*σ*_0_| < 1). Previous works have shown that this assumption has minimal implications for the resulting system behavior [40, 41]. The mean persistence time of the environment is *τ* generations (where a generation is defined as *J* duels); to model that, after each elementary time step the environment changes with probability 1*/Jτ*, and the fitnesses of all traits are re-drawn at random in an uncorrelated manner.

For simplicity we consider here only haploid genetic dynamics (i.e. asexual reproduction) but the model may describe allele frequencies in diploid populations provided that the genetic effect is additive.

Our model considers two genetic processes, mutation, generating new traits or subpopulation, and speciation, generating new species. Upon birth, an individual may produce a mutant offspring with probability *ν*. Such an offspring is the originator of a new *subpopulation*: this individual and its (nonmutant) descendants have a unique trait (or set of traits), so their response to the environment is independent of that of other sub-populations, still they belong to the same species. With a certain probability *µ* an offspring is the originator of a new *species* (and it picks a unique trait). Both mutations and speciation events are considered in an infinite-allele scenario, so each mutation or speciation yields a new subpopulation or a new species, and there are no recurrent mutations or recurrent speciations. Generally, the mutation rate *ν* governs the level of ITV, so we will focus on how this parameter affects all quantities of interest.

We will begin by analyzing the dynamics of this model with only two species, fixing *µ* at 0. In this case, one of the species will inevitably go extinct, and so the mean time to absorption, *T*_*A*_ quantifies coexistence. High *T*_*A*_ generally indicates that the system loses its diversity slowly, and therefore any factor or process increasing *T*_*A*_ promotes diversity. We would then consider the case of multiple species by taking *µ >* 0. In this case, we will study how species richness depends on mutation rate *ν*.

#### 2. Asymmetric model

To test whether our qualitative conclusions generalize to asymmetric (non-neutral) communities, we consider a high diversity community in a competitive system that allows for niche differentiation and fitness differences. A simple and popular model for such a community was proposed by [33, 34] and was calibrated for an empirical community of annual plants by Godoy and Levine [42]. In this model, the deterministic dynamics of species *i* is expressed as,

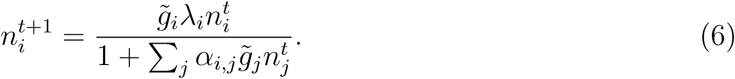

Here 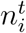 is the density of seeds of species *i* at time *t*, 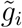 is the fraction of seeds that germinate and *λ*_*i*_ is the per-germinant fecundity of species *i* at low densities. The intra- and inter-specific interaction coefficients, *α*_*i,i*_ and *α*_*i,j*_ respectively, describe per-capita effect on seed production.

Following the analysis of similar models for competitive communities [43, 44] we choose *α*_*i,i*_ = 1 and pick all other (non-diagonal) elements of the competition matrix from a Gamma distribution with mean *C* and variance *γ*^2^. Accordingly, *C* measures the overall strength of competitive interactions (when *C* = 0 there is no inter-specific competition) and *γ* measures the heterogeneity of the competition matrix. The competition matrix elements *α*_*i,j*_ reflect niche overlap and fitness differences between species and are assumed to be time and trait independent.

Environmental variations are introduced through the germinant fecundity parameter which is taken to be time-dependent *λ*_*i*_(*t*). To emulate the effect of demographic stochasticity the fecundity of each germinant is drawn from a Poisson distribution with mean 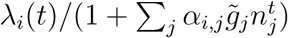.

Each population admits a genetic structure, so *n*_*i*_(*t*) = ∑*m*_*i,k*_(*t*) where *m*_*i,k*_ is the seed density of the *k*th subpopulation of the *i*th species. Different subpopulations have the same set of inter and intra specific competition coefficients but are subject to uncorrelated environmental variability, i.e, the parameters *λ*_*i,k*_(*t*) are IID, drawn from a uniform distribution between *J* (1 + *W/*2) and *J* (1 −*W/*2), where *J* sets the scale for community size and *W* is the amplitude of environmental fluctuations. Accordingly, for each subpopulation *k* the mean number of seeds in the next year is,

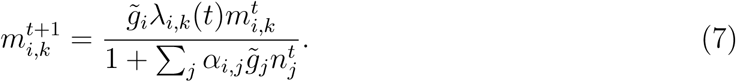

After each production-competition step (a year), each seed may mutate (i.e., becomes the originator of a new subpopulation) with probability *ν*. The system is subject to weak immigration from a regional pool with *S* species, where (like in [45]) the probability of immigration is a species independent constant (all species have the same density in the regional pool). In the simulation, each year one immigrant, drawn at random from the regional pool, arrives with probability *D*. We analyzed how the average species richness at equilibrium depends on mutation probability *ν* (which control the level of ITV in the system) at different parameter regimes.

## IV. RESULTS

### A. Quantifying the dynamical mechanisms

Let us consider a species with *N* individuals, which consists of M sub-populations of abundances *n*_*k*_, so *N* = *n*_1_ + *n*_2_ + …*n*_*M*_. If the fitness of the *k*th sub-population is *f*_*k*_(*t*), the average fitness of a randomly chosen individual in this population at time *t* is,

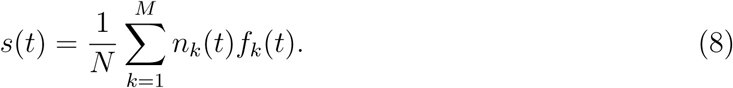

In an interspecific duel, since the chance to pick an individual from the *k*-th subspecies is *n*_*k*_*/N*, the mean fitness difference between two randomly chosen individuals, one belongs to species *i* and the other to species *j*, will be

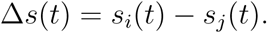

Accordingly, the chance of a species to gain or to lose an individual is determined by Δ*s*(*t*). This quantity itself fluctuates because of environmental variations and changes in subspecies composition.

A well known and widely used technique in the study of stochastic processes involves the diffusion approximation [19, 46, 47]. The essential aspects of this method are reviewed shortly in Supplementary I, where the derivation of Eq. 1 above is explained. The diffusion approximation characterizes all aspects of the dynamics, deterministic and stochastic, by the mean growth rate *s*_0_ and the variance *σ*^2^. Here we calculate numerically the mean of *s*_0_ and *σ*^2^ where expectation values are taken over histories and are denoted by 𝔼[·]. Accordingly, we use,

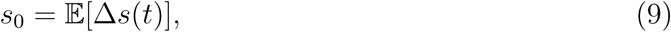

and

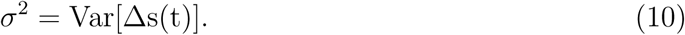

Note the distinction between *σ*_0_, which is the amplitude of variations of the fitness of a single **individual**, and *σ*, the amplitude of fitness variations of a given **species**.

Without ITV, *s*_0_ and *σ*^2^ are independent of the frequency of a species *x*. ITV generates frequency dependence, so one should consider the functions *s*_0_(*x*) and *σ*^2^(*x*). To demonstrate that, we ran simulations of the two species system with reflecting boundary conditions (i.e., upon extinction or fixation, the system was initiated again with *N* = 1 or *N* = *J -* 1, respectively). For any given *x*, the value of Δ*s* was monitored through time and its mean and variance were calculated.

The two mechanisms by which ITV affects ecological dynamics are manifested in the dependency of *s*_0_ and *σ* on *x*. In particular

- Due to the **adaptability advantage of diverse populations**, the mean fitness *s*_0_ of frequent species is higher than the mean fitness of rare species. Frequent species admits a standing phenotypic diversity so at every instance of time favoured sub-populations increase in frequency within the population, increasing the average fitness of a more variable species. Since the degree of ITV usually grows with abundance, *s*_0_(*x*) grows with *x* as exemplified in Figure 1. As a result, ITV acts to decrease the chance of invasion which is related to the growth rate when rare, *s*_0_(*x* ≪ 1). The dependence of this effect on *ν* is explained in detail in Supplementary II. When *ν* = 0, there is no ITV and no effect. As *ν* grows the resident species develops internal structure while the low-abundance invader is still homogenous, so the *s*_0_ of low-abundance species decreases with *ν*. For larger *ν* even small populations admit internal structure. Moreover, high fitness individuals often fail to inherit their traits to offspring due to high mutation rate. As a result, at too large *ν* the adaptability advantage of frequent species disappears.
- **Portfolio effect**. The effect of environmental variations on *s*(*t*) averages out when a species is composed of many independent sub-populations, so for a single population the variance of its fitness decreases with *x*. For two populations in a zero-sum game the variance of fitness differences, *σ*^2^(*x*) = 𝔼[*s*^2^(*x*)] + 𝔼[*s*^2^(1 −*x*)], has a minimum at *x* = 1*/*2.

This effect is demonstrated in Figure 2, where *σ*^2^ is plotted against *x* for different values of *ν*. Clearly, a species with richer internal structure is less sensitive to environmental variations, as these variations increase the fitness of some subpopulation and decrease fitness of others. Since the intra-diversity grows with the population size, large populations are less sensitive to environmental variations.

**FIG. 1:**
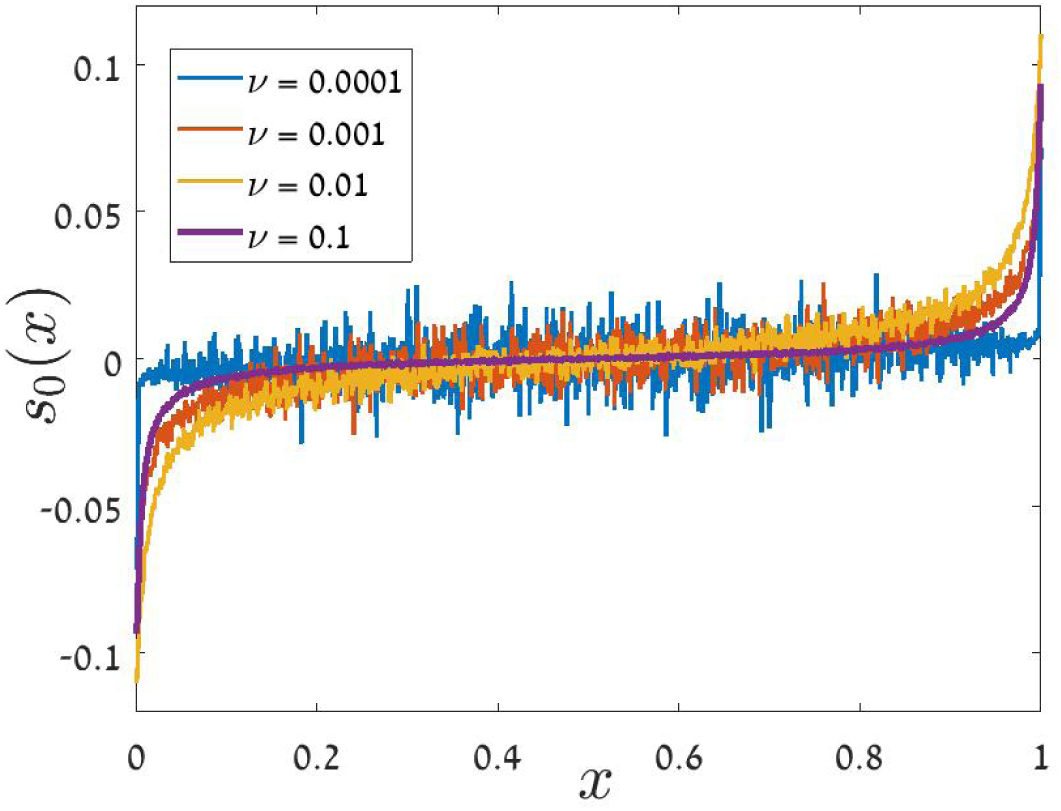
Adaptability advantage of diverse populations. The mean fitness *s*_0_(*x*) is plotted against the frequency *x*. We have simulated an individual based, two species competition model (*µ* = 0) with *J* = 1000, *σ*_0_ = 1*/*2 and *τ* = 1*/*4, and calculated *s*_0_(*x*) from (2). *s*_0_(*x*) is plotted for four different values of *ν*, see legend. A larger number of subpopulation is associated with higher fitness. Accordingly, low abundance species that have less structure are inferior. The relationship between *ν* and *s*_0_(*x*) is non-monotonic, see discussion in Supplementary II. The relative fitness of the rival species is just the minus of that of the focal speceis, hence the plot is anti-symmetric around *x* = 1*/*2.

**FIG. 2:**
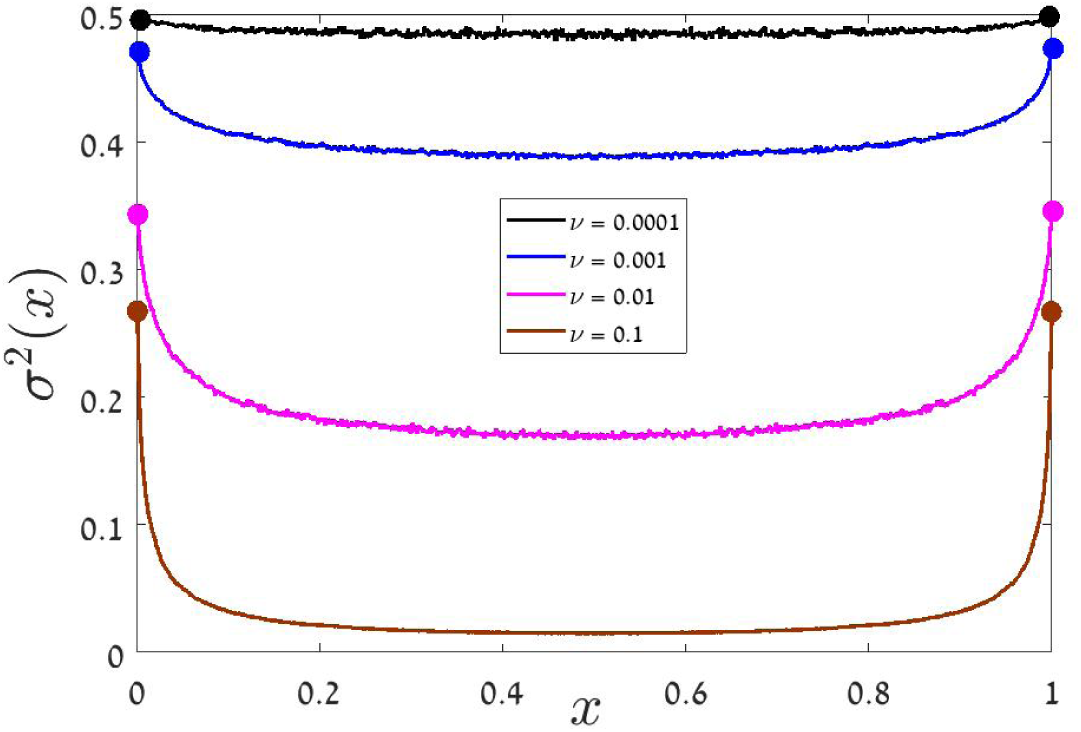
Portfolio effect. The variance of fitness variations, *σ*^2^(*x*) is plotted against the frequency *x* for two species competition with the same parameters used in Figure 1. *σ*^2^ has maxima close to the extinction/fixation points and minima at *x* = 1*/*2, where both species admit rich internal structure that buffers environmental fluctuations. The depth of the “smile” increases with *ν*, as expected. The first and the last point (*x* = 1*/J* and *x* = 1 − 1*/J*) of each dataset are marked by filled circle. Close to the extinction/fixation point, the minority species is almost homogenous. The majority species is diverse, and if *ν* is large its fitness is almost fixed. Accordingly, as *ν* increases *σ*^2^(1*/J*) is reduces to half of its value in a system without subpopulation structure.

Typically, at large *J* the portfolio effect turns out to be the dominant mechanism. To acquire the benefits of ITV - buffering against environmental variations and higher mean fitness due to a standing phenotypic diversity - a population must have *some* structure. Once this structure is obtained, the difference between a species with four subpopulations and one with twenty subpopulations is tiny. As a result, for fixed *ν*, a population of *N* individuals admits ITV with high probability when *νN >* 1. Accordingly, as discussed in Supplementary II, for a community of size *J* the resident species enjoys its higher adaptability against the invader only when *x* ≪ 1*/*(*νJ*), and this regime shrinks to zero for large community, i.e., when *J* → *∞*. On the other hand, the portfolio effect buffers a species against environmental variations whenever *x >* 1*/*(*νJ*), so this is the dominant effect in large communities.

### B. Two species competition model

In a previous study (28, see also Danino and Shnerb 27) the persistence of a two-species community (without ITV) was analyzed. Using the diffusion approximation technique illustrated in Supplementary I, Eq. (2) was derived for competition in temporally fluctuating environment and the validity of Eq. (1) for fixed environment has been discussed. In both cases the results depend only on *s*_0_ and *σ*^2^. Our two species model is simply a generalization of the dynamic considered by Danino et al., incorporating mutations that generate ITV.

In the previous subsection we showed how *s*_0_ and *σ*^2^ become frequency dependent in the presence of ITV and explained how this dependence is related to the portfolio effect and to the better adaptability of diverse populations. In this subsection we would like to demonstrate the effect of the these mechanisms on the persistence time of a community.

Both effects manifest themselves in the time to absorption (fixation or loss), *T*_*A*_(*x*) as demonstrated in Figure 3. For intermediate *x* values (the abundance of both species is not small) the portfolio effect dominates, increasing the time to absorption with respect to pure environmental stochasticity. As *ν* increases, the time to absorption approaches the purely demographic prediction (Eq. 1).

**FIG. 3:**
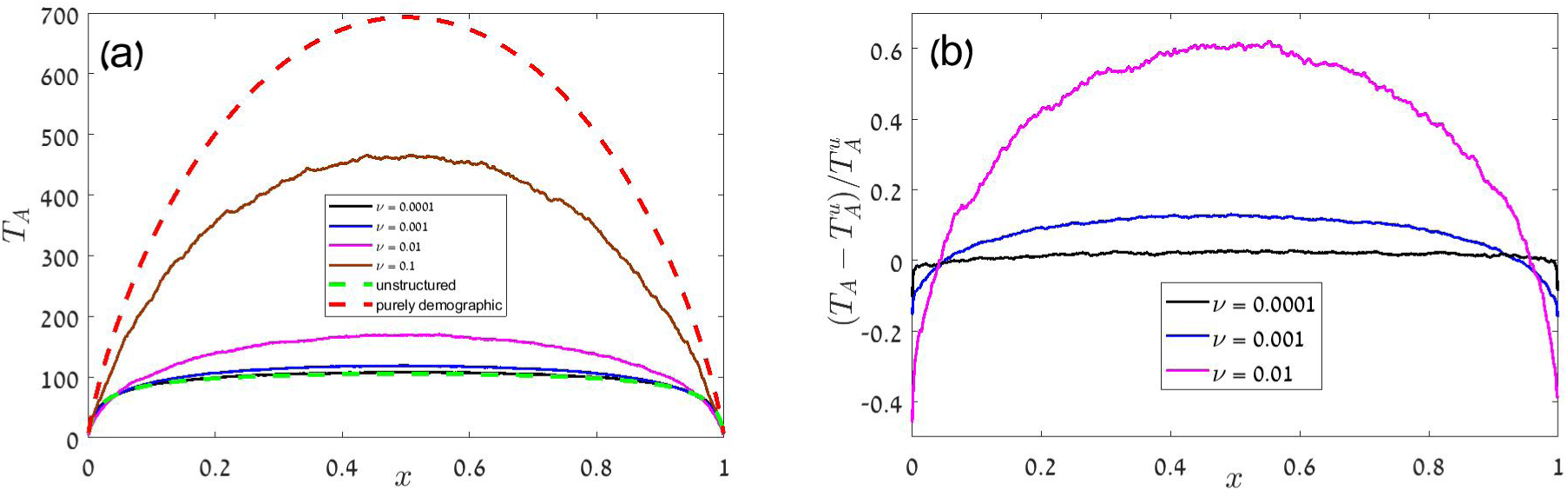
Time to absorption, *T*_*A*_(*x*), **vs**. *x* for two species competition with *σ*_0_ = 1*/*2, *J* = 1000 and *τ* = 1*/*4. In the left panel, the results for purely demographic stochasticity, 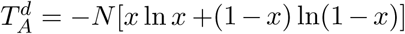 (dashed red line, Eq. 1), and for competition between two unstructured (no ITV) populations, (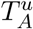, dashed green line, Eq. 2 with *g* = 2*σ*^2^*τ*) are compared with *T*_*A*_ for structured populations with different values of *ν* (full lines, see legend). Panel (a) demonstrates the portfolio effect: at higher *ν* the system becomes demographic like, as the effective value of *σ*^2^(*x*) decreases. In panel (b) 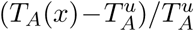 is plotted against *x* for the same datasets. The inferior adaptability of low abundance species causes 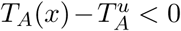 in the low-frequency regimes *x* ≪ 1 and at 1 −*x* ≪ 1.

To identify the effect of adaptability advantage one should focus on the small *x* region, as emphasized in Figure 3b. When a population becomes rare the advantage of the resident gets into play, decreasing *T*_*A*_ even below the prediction for purely environmental case (Eq. 2). As explained above, we expect that the frequency regime in which the adaptability effect dominates shrinks to zero as *J* increases.

### C. High diversity community - time averaged neutral model

The intuition gathered from the outcomes of two species numerical experiments suggest that in large communities the portfolio effect dominates intraspecific fitness advantage. Due to the portfolio effect the system becomes more and more “demographic like” as *ν* increases. In this section we would like to examine these phenomena for a community with many species.

Under pure demographic stochasticity and speciation events (*σ*_0_ = 0, *µ >* 0) one obtains the classical Kimura-Hubbel neutral model [38, 46, 48], where the species richness (SR) reflects an extinction-speciation equilibrium. The species abundance distribution has been shown to follow Fisher’s log-series statistics,

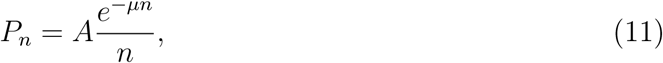

where *P*_*n*_ is the probability that a randomly picked species has abundance *n*. The normalization factor is approximately (when *µ*^2^ and exp(−*µJ*) terms are neglected) *A* ≈ (− ln *µ*)^−1^. When this approximation holds, the species richness is given by,

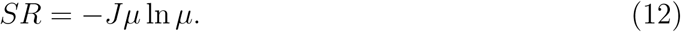

Recently, a neutral model with environmental fluctuations was suggested, and termed “time-averaged neutral theory” [9, 24, 32]. This model corresponds to our case of unstructured populations (where *ν* = 0 but *σ*_0_ *>* 0). The theory revealed that as *σ*_0_ increases, the species abundance distribution widens, decreasing evenness, so the total species richness decreases while the mean frequency of a species increases.

Figure 4 shows the effect of ITV on the species richness. As before, the two effects act in opposite direction. The lack of adaptability impedes the invasion of low abundance species so it leads to a decrease in species richness; this effect is seen to dominate the small *ν* sector. On the other hand the portfolio effect buffers against the environmental fluctuations so in the large *ν* sector the stochasticity becomes demographic-like and the richness increases.

**FIG. 4:**
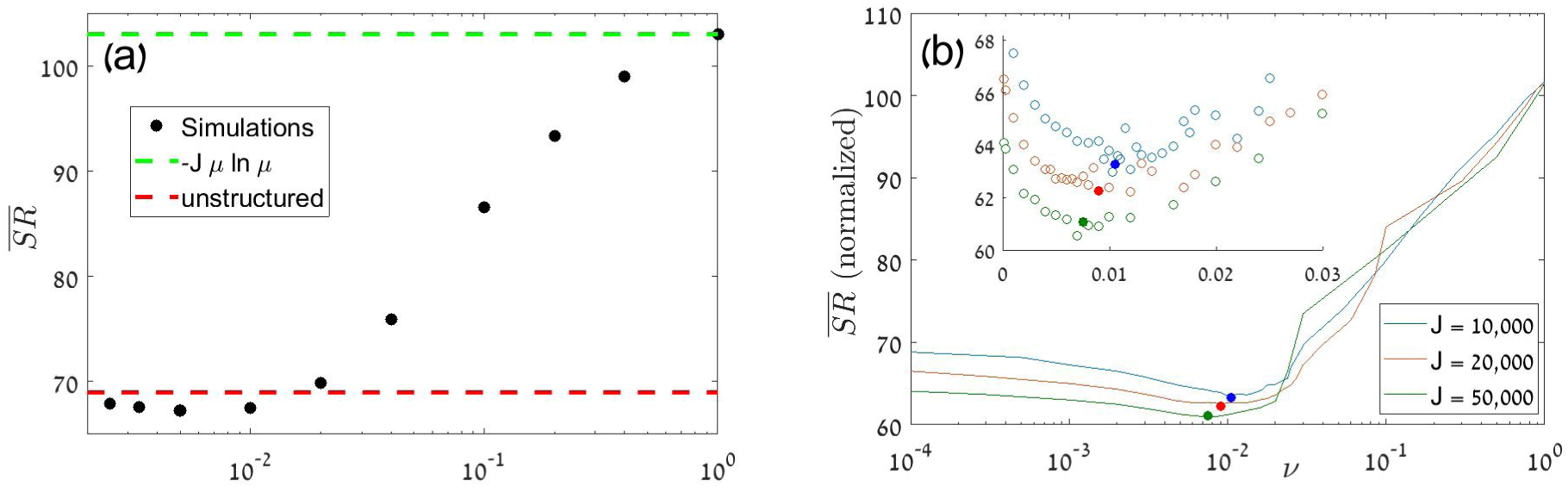
In panel (a), black dots shows the species richness 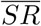 for a *J* = 10^4^ community with *σ*_0_ = 0.5, *τ* = 0.25 and *µ* = 0.0016, for different values of interspecific structure that arise from different values of *ν*. The prediction of Eq. (11) for pure demographic stochasticity is marked by dashed green line, and the mean species richness for a system with environmental stochasticity but without internal structure (*ν* = 0) is denoted by a red dashed line. An increase of *ν* first decreases the overall richness below the red line because of the inferior adaptability of low-abundance species, then it leads to an increase of 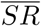 due to the portfolio effect, until it reaches the neutral limit. As explained in the text, we believe that the small *ν* sector (in which 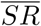 is reduced below its *ν* = 0 limit) disappears at large *J*. Our ability to simulate large systems is limited, however we can show that the minima is shifted systematically to lower values of *ν* as *J* increases. To that aim, panel (b) presents results for different values of *J* (when all other simulation parameters are the same). To allow for a decent comparison, in panel (b) 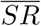 is divided by *J/*10000. The inset shows the non-smeared datapoints in the neighborhood of the minima (full circles).

As argued before, for any fixed frequency *x* the adaptability effect disappears and the portfolio effect strengthen as *J* increases, so we expect that the low *ν* sector disappears when *J* becomes large enough. Although we cannot demonstrate that completely, we present (in Fig. 4b) evidence for the narrowing of the low *ν* sector as *J* increases. Moreover, when we examined species abundance distributions (data not shown) it turns out that the reduction of 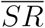 in the small *ν* sector is related to a reduction in the number of low abundance species that have inferior adaptivity, so the overall composition of the community is affected only weakly by this reduction and it will be quite difficult to observe the effect empirically.

### D. High diversity, asymmetric community - annual plant competition model

The asymmetric competitive Leslie-Gower model for annual plant community, as defined in section III A 2 above, provides us with another opportunity to test our predictions. In the absence of stochastic effects the factors that determine species richness are the number of species in the regional pool *S*, the typical level of niche overlap *C* and the typical level of fitness differences which is set by *γ*. The strength of demographic stochasticity is 1*/J* and the strength of environmental stochasticity is proportional to *W*, but its effective value decreases with *ν*.

Figure 5 shows species richness, as measured numerically, as a function of *ν* for different values of *C, γ* and *W*. In all cases the species richness increases with *ν*, since an increase in *ν* diminishes the negative effect of environmental stochasticity on species richness. Note that for such a model, with non-overlapping generations and uncorrelated environment, there is no adaptability effect since the environmental conditions change after each elementary event of intraspecific competition.

**FIG. 5:**
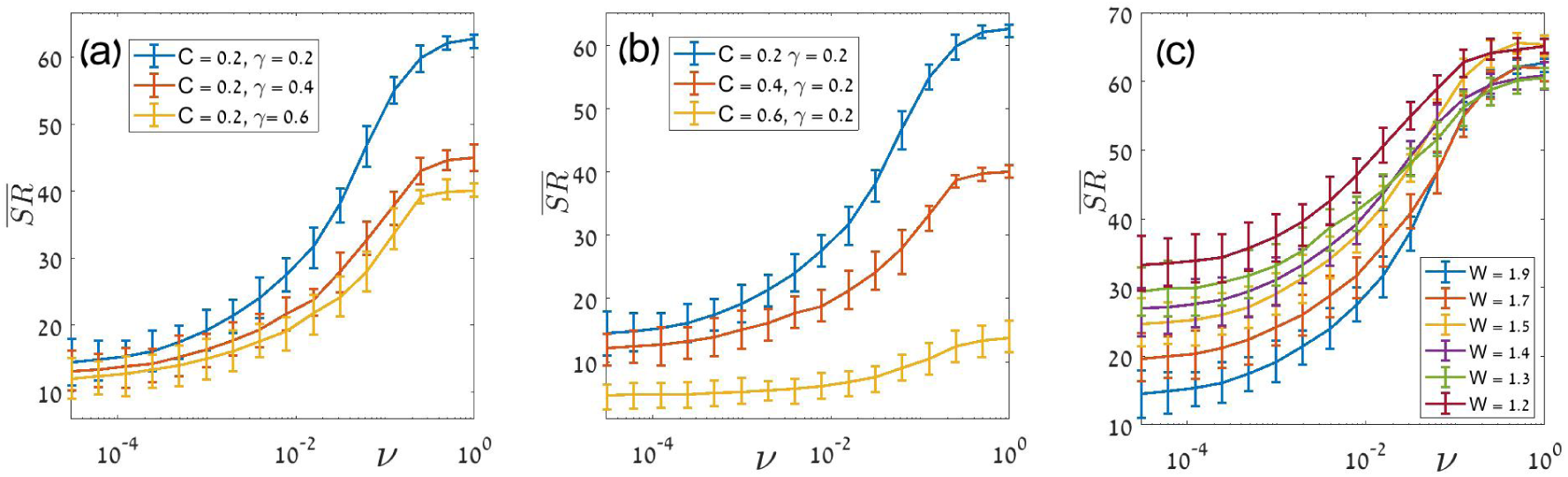
Species richness for a competitive Leslie-Gower model for annual plant community, where the regional pool has *S* = 70 species. The non-diagonal terms of the competition matrix *α* were drawn independently from a Gamma distribution with mean *C* and variance *γ*. For each subpopulation at each year, *λ* was drawn from a uniform distribution between *J* (1 + *W/*2) and *J* (1 −*W/*2), where *J* = 10000. For all subpopulations 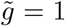. The system was initiated with all *S* (genetically homogenous) species at equal densities *J/S* and results were collected (after 2000 equilibration steps) through 400000 consecutive years. In each year one immigrant, drawn at random from the regional pool, arrives with probability 0.1 (a single immigration event every 10 years, on average). In panel (a), species richness at equilibrium is plotted against *ν* for *W* = 1.9 and *C* = 0.2 for three values of *γ*. In panel (b), *W* = 1.9 and *γ* = 0.2, *C* = 0.2, 0.4 and 0.6, while in panel (c) *C* = 0.2, *γ* = 0.2 and different values of *W* were used. In all cases species richness increases with *ν*. Error bars represent the interval between the 5th percentile and the 95th percentile of the *SR* distribution.

## V. DISCUSSION

In this work we have identified two mechanisms by which ITV influences species diversity in a fluctuating environment. First, ITV breaks the synchrony between individuals in the population, buffering fluctuations in population size and effectively partially transforming environmental fluctuations into demographic-like stochasticity. The second mechanism is the adaptability advantage of diverse populations, where the fitter individuals reproduce within the population, increasing the fitness of a diverse species relative to more homogeneous populations. These two mechanisms act in opposite directions on species diversity. The portfolio effect limits the magnitude of population fluctuations and increases the time it takes populations to go stochastically extinct. On the other hand, since invading species typically show low ITV, the adaptability effect reduces their relative fitness, making invasion harder and diversity lower. We further studied the net effect on species diversity in two- and multi-species communities. We found that the portfolio effect is dominant for large communities, considerably increasing species richness in multiple cases, while the potential reduction in diversity due to the lack of adaptability of rare populations becomes weaker as the size of the community increases.

To study the portfolio effect and the adaptability advantage we used very simple models to attain generality, finding that the portfolio effect is stronger in large communities, leading to a net positive effect of ITV on species diversity. We believe that these mechanisms should generally play some role in every community. Yet, we ignored the deterministic effects that ITV may have in a fixed environment on fitness and niche differences, as these have been studied elsewhere [15, 16]. It is possible that these effects of ITV would interact with the mechanisms we identified, and the net effect of ITV would depend on the multiple ways that ITV influences the multiple aspects of community dynamics. More work is needed to identify how the overall effect of ITV on species diversity is determined.

We considered a simple eco-evolutionary dynamics with perfect heritability (when mutation does not occur) and completely uncorrelated response of different traits to the environment.In reality one expects more gradual process in which intra-specific trait differences reflect the accumulation of many elementary genetic mutations. However, even in realistic situations with many types of environmental variations, partial heritability and gradual trait variability, the dynamics of a population depends on the response of the mean and the variance of its fitness to a given level of environmental fluctuations. Therefore we believe that one may take complex situations into account using an effective mutation rate. Another possibility would be to take into account the covariance between different individuals when *σ*^2^ is calculated.

We further assumed a mode of competition that (in the absence of ITV) does not lead by itself to a disadvantage or advantage of rare species (e.g. storage effects, [8, 9]). Without storage, an increase in the amplitude of temporal environmental fluctuations has only negative effect on species richness, and this allows us to identify the influence of the portfolio effect unambiguously. In the presence of storage and storage-like effects, an increase of the amplitude of environmental fluctuations have contrasting consequences: on the one hand, it increases the invasion of rare species, but simultaneously it enhances stochastic extinctions and lower diversity [9, 32]. Hence, the net effect of environmental fluctuations on species-diversity is not trivial. For this reason, it is difficult to evaluate the effect of ITV that influences the response to such fluctuations, and it will likely depend on the specific parameters. Yet, since ITV has the general effect of transforming environmental fluctuations into demographic-like stochasticity, we hypothesize that ITV will weaken both the positive and negative effects of the storage mechanism. More work is required to develop a synthetic understanding of the effect of ITV on species diversity in fluctuating environments that would consider more realistic and complex scenarios.

From a broader perspective, species richness is a result of the balance between colonization and extinction processes [49]. Yet, in recent years there is an increasing interest in analyzing species diversity using modern coexistence theory, which considers growth rate when rare as the measure for the ability of species to coexist and enhances diversity [2, 50]. Our result reveal the contrast between these two views, as ITV reduces the growth rate of rare species, impeding deterministic coexistence, but its net effect on species diversity is often positive because it also reduced the magnitude of stochastic extinctions. We believe that this is a general feature, and whenever a mechanism has both deterministic effects and stochastic effects, using modern coexistence theory to understand its effect on diversity may reveal only part of the picture, and not necessarily the dominant part. Apparently, studying the time to extinction of populations may in many cases be more revealing [11].

## VI. ACKNOWLEDGMENTS

This research was supported by the ISF-NRF Singapore joint research program (grant number 2669/17).

## Supplementary information

### I. THE DIFFUSION APPROXIMATION

The diffusion approximation is a popular technique that allows one to analyze the dynamics of stochastic systems, even when the stochasticity is superimposed on deterministic forces. The technique is widely used in population genetic [35, 47] and theoretical ecology [19] and is known to be remarkably robust to modifications of the microscopic dynamics of the model [51].

Let us give an example, using a simplified version of the model defined in the main text. In two species competition, when the frequency of the focal species is *x* = *n/J* (*n* is the number of individuals, *J* is the total size of the community), each elementary birth-death event involves interspecific competition with probability 2*x*(1 −*x*). By definition the time needed for an elementary step is 1*/J* generations. If the dynamics is completely *neutral*, in every interspecific competition event the focal species gains one individual (*x* → *x* + 1*/J*) with probability 1*/*2 and loses one individual (*x* → *x* −1*/J*) with probability 1*/*2.

To calculate the mean time to absorption (either fixation or loss) for a system with *n* individuals, *T*_*A*_(*n*), one should consider all possible states available for the system after a single elementary event, in our case these are *n* and *n* ± 1. The mean time *T*_*A*_(*n*) is obtained as the time required for a single step, plus the sum of the mean time from all possible destinations, weighted by the chance to reach the destination state. Accordingly, in the neutral case the mean time to extinction satisfies the difference equation,

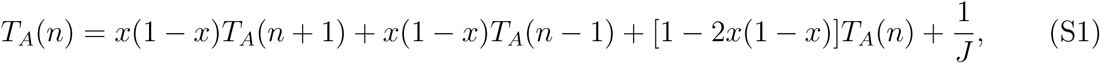

with the boundary conditions *T* (0) = *T* (*J*) = 0.

Equations like (S1) are usually difficult to solve. The essence of the diffusion approximation is to replace *n* by *x* and to expand each function of *x* ± 1*/J* to second order in 1*/J*, so 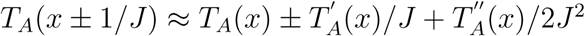. Plugging that into (S1) one finds,

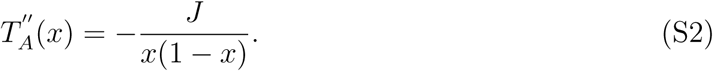

The solution for this equation with the boundary conditions *T* (0) = *T* (1) = 0 yields Eq. (1) of the main text.

In general [as explained in detail in textbooks like [47]], to implement the diffusion approximation one calculates the mean growth rate at a certain frequency, *s*_0_(*x*), and the corresponding variance *σ*^2^(*x*), and plugs these quantities into the backward Kolmogorov equation,

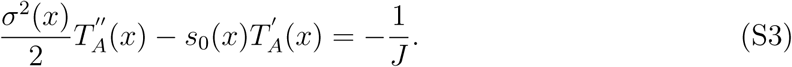

Eq. (S2) is a special case of Eq. (S3) where *s*_0_ = 0 (there is no directionality since the model is neutral) and *σ*^2^ = 2*x*(1 −*x*)*/J* ^2^. This quantity is indeed the variance when the chance, for both 1*/J* gain and 1*/J* loss, is *x*(1 −*x*).

The implementation of the diffusion approximation is based on the assumption that the function *T*_*A*_(*n*) is smooth enough over the integers, such that *T* (*n* + 1) may be approximated by the first and the second derivative evaluated at *x* = *n/J*. As discussed in length in [28], this approximation fits perfectly the numerical results in the purely neutral case (Eq. 1 of the main text) and for two species under demographic and environmental stochasticity but without internal structure (Eq. 2 of the main text).

### II. FEATURES OF *s*(*x*) AND *σ*^2^(*x*)

In the main text we have identified two mechanisms that affect the dynamics of a genetically structured population, the excess adaptivity of abundant species due to their internal structure and the portfolio effect that buffers the response of the mean fitness of composite population against environmental variations.

These effects were demonstrated, for a few specific systems, in Figures 1 and 2, where *σ*^2^(*x*) and *s*_0_(*x*) were plotted against *x*. In these figures we have used systems with specific values of *J*, *ν, τ* and *σ*_0_. Here we would like to discuss the general dependencies of *σ*^2^ and *s*_0_ on these parameters.

Let us begin with the decrease of *s*_0_ with abundance, as depicted in Figure 1. In fact, the mean fitness of small populations decreases for two reasons. One, that has nothing to do with ITV, is that populations usually become rare when their fitness is low. The other reason is the growth of fitter subspecies that increases the overall fitness of composite species, so small populations that are usually homogeneous become inferior. Both effects peak for a singleton, so a rough measure of the overall covariance between abundance and fitness is given by *s*_0_(1*/J*).

Figure S1 shows how the mean fitness of singletons varies with *ν*. When *ν* = 0 (no ITV) a singleton is still inferior because of the first reason. As *ν* increases ITV contributes to the effect and *s*_0_(1*/J*) decreases even further. However, above some value of *ν* the effect shrinks substantially, as mutations inhibit the growth of temporaly superior subpopulations.

**FIG. S1:**
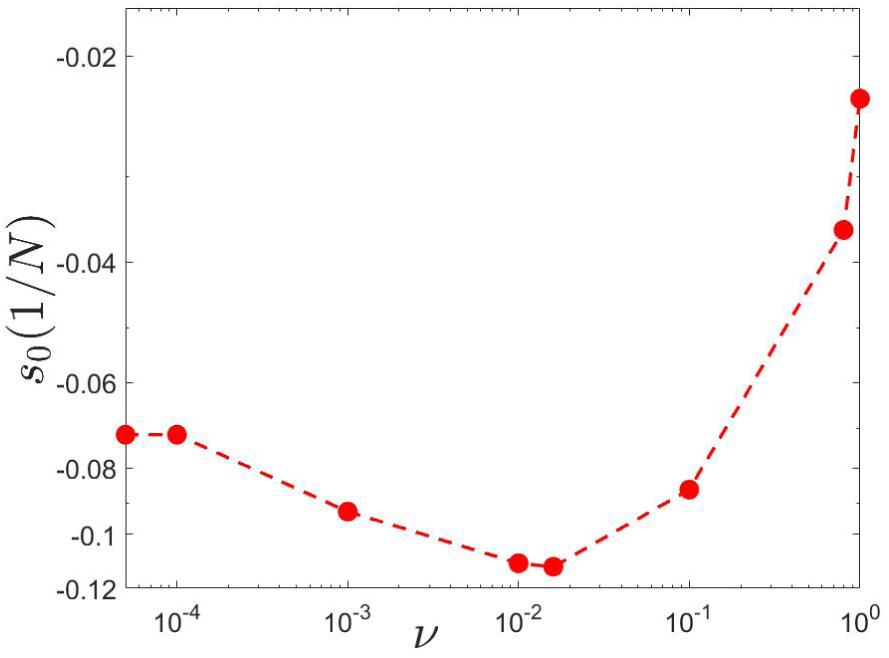
Covariance between abundance and fitness, as manifested in *s*_0_(1*/J*), is plotted against *ν* for *J* = 1000, *σ*_0_ = 0.5 and *τ* = 0.25. Black circles are the outcomes of numerical simulations, the dashed line was plotted to guide the eye.

As already seen in Figure 1, the excess adaptivity of the abundant species manifests itself only close to the extinction/fixation points. To qualify for the benefit of higher adaptivity, only a few subpopulations are needed. As a result, one may expect that for populations with *n*_*c*_ *> c*_2_*/ν* (where *c*_2_ is some order one constant) the excess adaptivity of the abundant species disappears. The value of *n*_*c*_ is independent of *J* as demonstrated in Figure S2. This implies that the above frequency *x*_*c*_ = *n*_*c*_*/J* the selective disadvantage of small populations disappears, where *x*_*c*_ → 0 as *J* increases.

The portfolio effect manifests itself in the dependence of *σ*^2^ on *x*. Without internal structure each species has fitness *σ*_0_ with probability 1/2 and −*σ*_0_ with probability 1/2, so the fitness difference is either zero or *±*2*σ*_0_ and the variance is 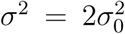. This was demonstrated in Figure 2 above, where for small *ν* (almost homogenous populations) *σ*^2^ is close to 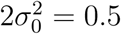.

**FIG. S2:**
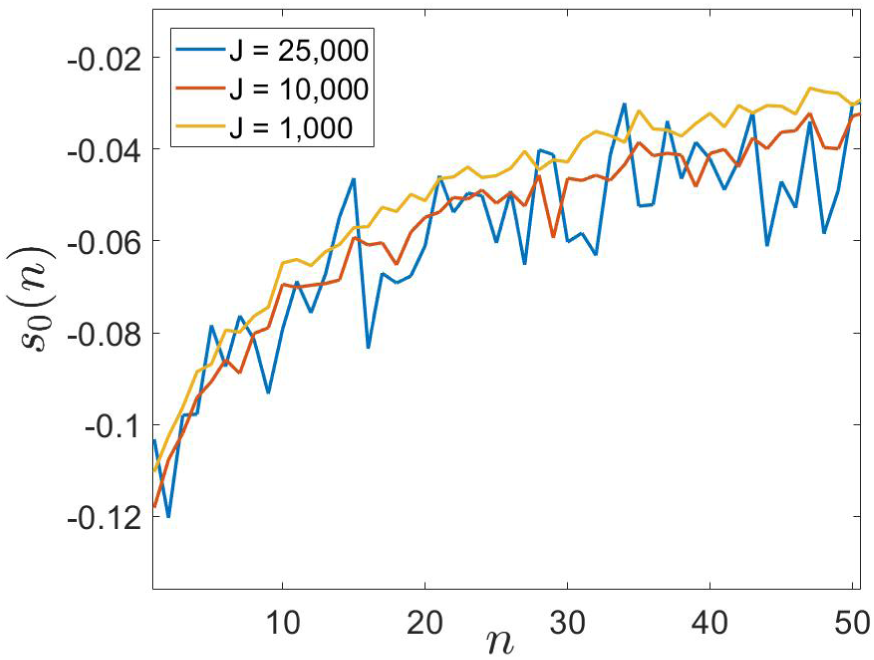
*s*_0_(*n* = *xJ*), is plotted against *xJ* for different values of *J* (see legend). *xJ* is the number of individuals of the focal species, and one sees that the effect depends only on absolute number, not on frequency. Accordingly, the frequency regime over which the abundant species is superior shrinks like 1*/J*. Results were obtained from MC simulations with *ν* = 0.01, *σ*_0_ = 0.5 and *τ* = 0.25.

Clearly, *σ*^2^(*x*) has a minimum in *x* = 1*/*2, where both species admit rich subpopulation structure. In that case, the large *J* behavior of *σ*(1*/*2) strongly depends on the ratio between environmental stochasticity and the rate of mutation.

As shown in [24], as long as *ν >* 2*g* (where *g* = *σ*^2^*τ/*2) subpopulations with abundance above 1*/*(*ν -* 2*g*) are exponentially rare. This implies that all the subpopulations are “microscopic” - their abundance does not scale with *J*. Accordingly, when *J* → *∞* the number of subpopulations at each abundance class is diverging, and since the fitness of each subpopulation fluctuates independently, *σ*^2^(1*/*2) → 0.

On the other hand, when *g > ν/*2 the abundance of some subpopulations is a finite fraction of *J* (say, *J/*10), so as *J* grows the number of these “macroscopic” subpopulations is kept, more or less, fixed. These subpopulations grow in abundance with *J* and their contribution to *σ*^2^ does not averages out. As a result, *σ*^2^(1*/*2) approaches a finite value even in the *J* → *∞* limit.

